# A plea for evidence in ecosystem service science: a framework and its application

**DOI:** 10.1101/007021

**Authors:** Anne-Christine Mupepele, Carsten F. Dormann

## Abstract

The ecosystem service concept is at the interface of ecology, economics and politcs, with scientific results rapidly translated into management or political action. This emphasises the importance of **reliable recommendations** provided by scientist. We propose to use evidence-based practice in ecosystem service science in order to evaluate and improve the reliability of scientific statements. For this purpose, we introduce a level-of-evidence scale ranking study designs (e.g. review, case-control, descriptive) in combination with a study quality checklist. For illustration, the concept was directly applied to 12 case studies. We also review criticisms levered against evidence-based practice and how it applies to ecosystem services science. We further discuss who should use the evidence-based concept and suggest important next steps, with a focus on the development of guidelines for methods used in ecosystem service assessments.

---

Ecosystem services, the benefits humans derive from nature, have gained popularity over the past ten years (Raffaelli and White, 2013). The concept provides a common discussion ground in science-policy interaction (Daily *et al*., 2009). Beside the positive aspects of increasing popularity and public attention, it runs the risk to serve as a buzzword boosting scientifically weak studies (Vihervaara *et al*., 2010). To lend scientific credibility to the ecosystem services concept, we need to improve the scientific basis of ecosystem services, together with an increased awareness about the reliability of current results (Carpenter *et al*., 2009; Boyd, 2013).

It was medicine that pioneered the evidence-based concept assessing the reliability of scientific statements and encouraging practitioners (doctors) to use only the most solid recommendations (Sackett *et al*., 1996, Cochrane Collaboration - www.cochrane.org). In evidence-based medicine, scientific results are ranked hierarchically according to their study design and quality (OCEBM Levels of Evidence Working Group, 2011). Such a scale permits the identification of the most reliable recommendation for diagnoses and treatments.

New concepts entail evaluation, and evidence-based practice has not stayed without criticism. We discuss the central arguments raised against evidence-based practice. Despite this criticism, evidence-based practice is successfully implemented and applied in medicine, today. The concept is also mentioned in other areas, including justice (www.campbellcollaboration.org), economics (Reiss, 2004) and environmental science such as conservation (Pullin and Knight, 2001, 2009; Sutherland *et al*., 2004) or forestry (Binkley and Menyailo, 2005; Petrokofsky *et al*., 2011).

In environmental science the most relevant step towards an evidence-based practice were the introductions of the journals ‘Conservation Evidence’ in 2004 and ‘Environmental Evidence’ in 2011, by the Collaboration for Environmental Evidence (www.environmentalevidence.org). The editors were the first to transfer evidence-based medicine to conservation (Pullin and Knight, 2001). Discussions arose about the hierarchy of study designs that should be used in environmental science. Pullin and Knight (2001) and Petrokofsky *et al*. (2011) encouraged the use of a scale closely related to medicine, but this scale did not represent well the approaches normally used in environmental science. Sutherland *et al*. (2004) argue that we cannot use a hierarchy at all because conservation, and environmental science more generally, is less straightforward and less well resourced than medicine. Nevertheless these authors agree that the top of the hierarchy, the gold standard, is represented by systematic reviews, and therefore the Collaboration for Environmental Evidence highly emphasises the generation of systematic reviews (Pullin and Knight, 2009; Sutherland *et al*., 2004; Petrokofsky *et al*., 2011). Systematic reviews are not the only source of information for practitioners, scientists and policy makers and evidence-based practice involves tracking down the best available evidence with which to answer the question at hand (Sackett *et al*., 1996).

Our aim is to propose a hierarchy and a quality checklist ranking the strength of evidence of common study designs in combination with quality criteria. These are valid for all environmental science studies. We further introduce evidence-based practice to ecosystem service science, which has not yet seen it in use. Scientists and decision makers should elucidate and transparently quantify the reliability of knowledge and thus the scientific basis for decisions taken. We give clear guidance on the terminology around evidence-based practice, to ensure that scientists and practitioner can communicate effectively across the disciplines and backgrounds. In the last section we provide examples for the application of the concept, respond to common criticism and offer suggestions for the next steps.

## The evidence-based concept

The terminology used around evidence-based practice is diverse even in the medical field. However a well-defined terminology is essential for good communication between practitioners and scientists.

According to the Oxford English Dictionary, evidence is the available body of facts or information indicating whether a belief or proposition is true or valid (www.oxforddictionaries.com/definition/english/evidence). In other words, **evidence** is a measure for the knowledge behind a statement. The **strength of evidence** reflects the quality of our knowledge and we can identify whether a statement is based on **high or low evidence**, hence very reliable or hardly reliable. Following this argumentation, **evidence-based** practice means that actions are undertaken based on knowledge of the highest available reliability. It further means that if high evidence results are missing, the end-user is aware about the low reliability of the statement.

Evidence-based practice starts with a question or a purpose (Fig. 1). The way to the answer, i.e. to the outcome of the study, implies a study design. The study design is the set-up of the investigation, controlled, correlative or observational. These study designs are not equally good, leading to different strengths of evidence. In order to derive a level of evidence, we need a hierarchical scale ranking study designs. Further the implementation of the design is important, and assessed in the critical appraisal. Study designs with a high level of evidence can be implemented poorly. We provide a quality checklist to derive the study quality further below. With help of the critical appraisal we determine the final level of evidence, depending on the study design as well as on quality criteria.

**Figure 1.**
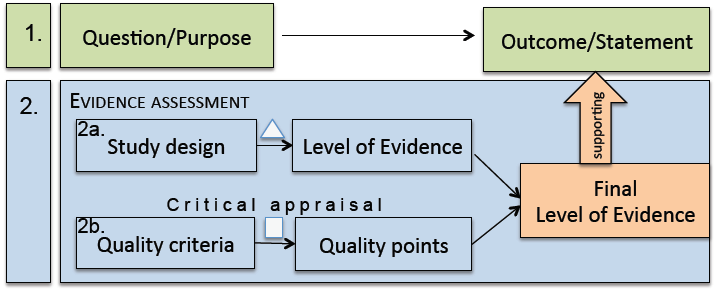
Schematic procedure in evidence-based practice: 1. Identification of question/purpose of the study and the outcome/statement, given as result of the study. 2. The assessment of the evidence supporting the outcome, with help of a level-of-evidence pyramid (Δ) and a quality checklist (□).

### 1. Question, outcome and the context

As in all of science the purpose of the investigation, ideally in form of a question, has to be clear. Still, it is sometimes surprisingly challenging to ask a question correctly. For example, the question has to fulfil certain criteria to be a well-focused and must be an answerable question (Higgins and Green, 2011; Collaboration for Environmental Evidence, 2013, p.20-23). For ecosystem service science, we suggest in addition to the question the specification of the environment and the context. The information *which* ecosystem service is investigated in *which* system is necessary to determine the context for the validity of the answer. Ecosystem service science is interdisciplinary and combines ecology, economy, politic and other social and natural sciences. In order to know which field we operate in, it is recommended to determine the facet of the ecosystem services question:

1. **Quantification** of ecosystem services: the amount of an ecosystem service or a set of services. It can be measured in absolute units or relative to another system.
2. **Valuation** of ecosystem services: the societal value of a service or a set of services. The most common way is monetary valuation. Other possibilities are in relation to a reference system or on a ranked scale (high, middle, low value).
3. **Management** of ecosystem services: the management/treatment of an ecosystem to favour specific ecosystem services. For example leaving dead wood in forests to increase biodiversity or reducing agricultural fertiliser to decrease nearby lake eutrophication.
4. **Governance** of ecosystem services: the strategy to steer a management type. The tools used are either incentives (subsidiaries) or penalties (law/tax).

Ideally these facets are investigated in the presented order starting with the quantification of an ecosystem service, which should then be valued. The most valuable services will be favoured by a well-adapted management option and in the end a governance strategy of how to steer the preferred type of management is implemented. Deviations of this structure are common, e.g. valuation does not necessarily require prior quantification. However, to cover the whole width of ecosystem service science, all four steps are required.

We have highlighted the question, context and facet. In an ecosystem services study, this is followed by the actual investigation. The outcome is usually the result of the study, it is the answer to the originally formulated question.

### 2. Evidence assessment

The outcome of an investigation can be of high or low reliability depending what was done to achieve the answer. The evidence assessment investigates the study design and the quality in order to determine the reliability of the outcome. In the following we present an evidence assessment not only for ecosystem service science, but also for all other environmental sciences.

#### Level-of-evidence pyramid

At the heart of evidence-based practice lies the hierarchy to rank the study designs (Fig. 2). The study design determines whether it yields high or low evidence. **Systematic reviews (LoE1a)** are at the top end of the level-of-evidence scale and provide the most reliable information. They summarise all information gained in several individual studies and are conducted according to strict guidelines (e.g. Collaboration for Environmental Evidence, 2013). Ideally they include quantitative measures, at best a meta-analysis (in the strict sense; see Borenstein *et al*., 2009; Vetter *et al*., 2013). Other more **conventional reviews (LoE1b)** may also include quantitative analysis or be purely qualitative. They both summarise the findings of several studies, but conventional reviews are less complete, not reproducible and often suffer more from publication bias.

**Figure 2.**
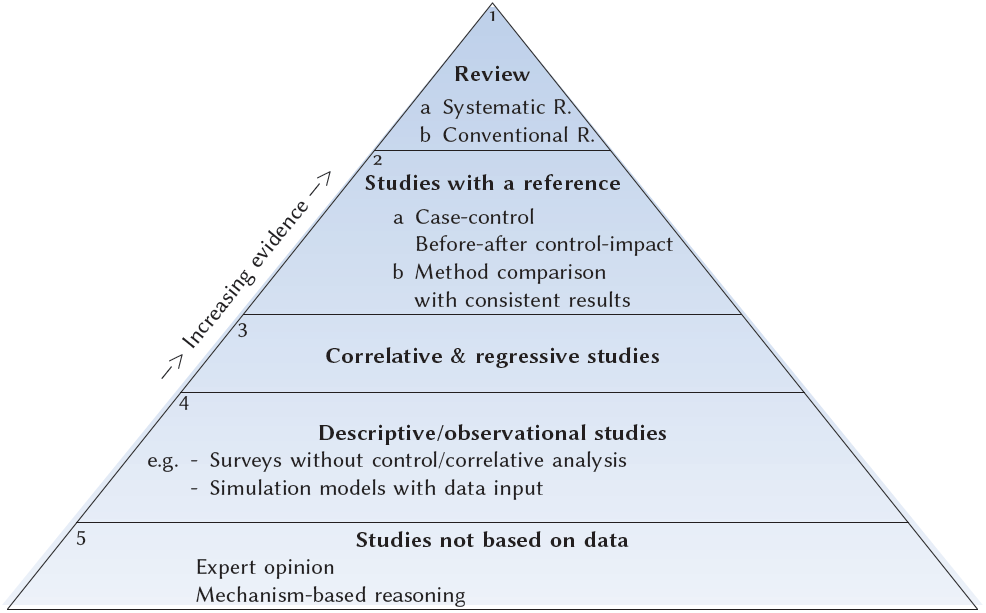
Level-of-evidence (LoE) pyramid ranking study designs according to their evidence. LoE1 - LoE5 with subcategories a and b.

The necessary condition for any review is that appropriate individual studies are available. The most reliable individual studies are **studies with a reference (LoE2)**. Typically, these are case-control or before-after control-impact studies. Method comparison can be useful for the valuation of ecosystem services, where no ‘true’ reference exists, however the results between both methods have to be consistent to provide high evidence.

Uncontrolled **correlative and regressive studies (LoE3)** are studies investigating for examples the influence of environmental variables on the quantity of an ecosystem service. **Descriptive studies, also called observational studies (LoE4)** present the data collected, sometimes in summary statistics or ordinations or they feed into simulation models. They are based on data, but not conducted in a controlled or correlative design.

The lowest level of evidence are statements that are **not based on any data (LoEs)**. These are usually anecdotes or expert opinions, the latter ones often not better than random (Tetlock, 2005). Even if their argumentation is a mechanism-based reasoning (‘first principles’: *A* works according to a certain mechanism, so we expect *B* to work in the same way), we cannot rely on these statements in the context of ecosystem services, where no first principles exist (Lawton, 1999).

It is important to note that ‘method’ and ‘design’ should not be confused. Methods are the means used to collect or analyse data, e.g. remote sensing, questionnaires, ordination techniques, model types. The design reflects how the study was planed and conducted, e.g. a case-control or descriptive design. For some methods, the underlying design is not easy to identify. Remote sensing for example can be done purely descriptive or with a valid reference such as ground-truthing or in a ‘before-after’ design. Most methods used in a descriptive design could actually follow a controlled design, but not necessarily do so.

#### Critical appraisal

The critical appraisal assesses the quality of the implementation of a study design. A study with a high evidence design may be poorly conducted. The critical appraisal identifies the study and reporting quality. It may lead to a correction of the level of evidence, so that the final level of evidence supporting the outcome is lower than the one allocated according to the design. This depends on objective, sometimes design- or facet-specific criteria. Several literature sources provide lists with quality criteria (e.g. Rychetnik *et al*., 2001; Pullin and Knight, 2003; Söderqvist and Soutukorva, 2006; Balshem *et al*., 2011, Oxford Centre for Evidence-based medicine 2011 www.cebm.net). We combined these lists to a general quality checklist (Box 1). The checklist consists of 33 questions with the possibility to use only a subset if some questions are not appropriate for the specific context. All questions answered with yes receive one point (or two points if it is an important questions - in bold font in Box 1), and zero points if answered with no. In case of non-reported issues, we advice the answer ‘no’ to indicate a deficient reporting quality. The percentage of points received out of possible points will help to decide whether to downgrade the level of evidence.

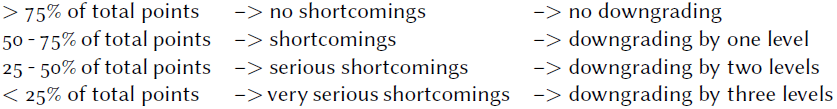

**Box 1.**
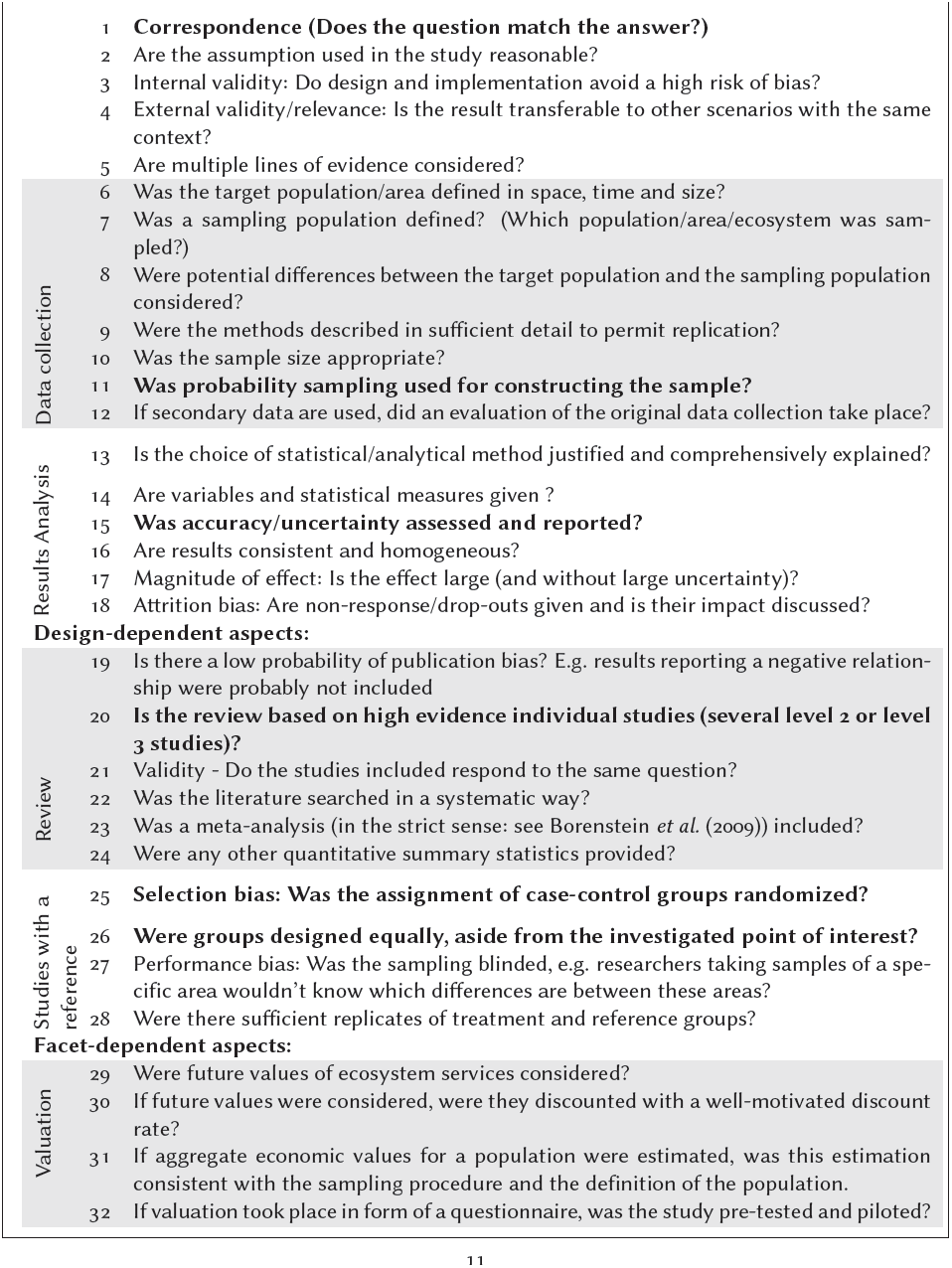
The **quality checklist** is designed in form of questions. Each question answered with ‘yes’ will receive a point, important aspects (bold type) two points. If a question is not appropriate in the specific context, it may be left out.

For example, if the first 17 questions of the checklist (Box 1) were answered, 10 of them - including the 3 bold ones - with ‘yes’ and 7 with ‘no’. 13 out of 20 points (65%) were reached. 65% means that there are shortcomings and it is suggested to downgrade the study by one level of evidence.

We encourage the use of the checklist for an orientation, but we want to emphasise that this procedure can not be fully standardised. Quality aspects can also depend on the context of the study and the final judgement will remain with the user. Reviews provide information on the highest level of evidence and the critical appraisal is different from other designs, because they themselves are based on studies with lower evidence (see Box 1: section review). If only studies based on low evidence were included, the quality assessment should downgrade a review to LoE4 and if in addition other quality issues showed serious shortcomings even to LoE5.

## Application of the evidence-based concept

The most popular application of the evidence-based concept is a systematic review that is used to summarise all knowledge available for a specific question. A systematic review is however time consuming and if policy makers need a specific answer in a shorter time, a ‘rapid evidence assessment’ (UK Civilservice, 2013) can be used as an alternative to a systematic review. Another approach to evidence-based practice are synopses. Synopses do not focus on a specific question but bring together information from a much broader topic, e.g. from a whole animal class, such as amphibians (Smith and Sutherland, 2014). A third possibility to use the evidence-based concept are guidelines to recommend tools/methods based on the best available evidence. These ‘best practice guides’ will focus on methods and the questions are therefore less typical systematic review questions, e.g. ‘How much CO_2_ is stored in European temperate forests?’, but more like ‘Which is the best method to measure CO_2_ stored in temperate forests?’ This serves to allow forest scientists to employ the best method to any temperate forest. In the case of evidence-based ecosystem service science that would also identify the evidence base of common instruments and tools, e.g. INVEST (Tallis and Polasky, 2009). All these possibilities for the application of the evidence-based concept summarise individual studies and therefore require the evaluation of the evidence of individual studies included. In systematic reviews this is typically done as a step in the critical appraisal, but so far a scale and a clear guideline was missing. With the method described above we can assess the level of evidence of individual studies and in the following we provide several examples (more details in the supplement table S1 and S2).

### Examples of evidence-based practice

**‘**How does adding dead wood influence the provision of ecosystem services?’ was a question addressed by Acuña *et al*. (2013). They investigated two ecosystem services (food (fish) and retention of organic and inorganic matter) in a river-forest ecosystem in Spain and Portugal and studied the effect of a management intervention. Their study design followed a before-after control-impact approach, which is LoE2. The critical appraisal (see supplement table S2) revealed shortcomings: only 14 out of 24 points (58%) were gained. The level of evidence was downgraded by one level to level three. We therefore conclude that the statement made by Acuña *et al*. (2013): ‘restoration of natural wood loading in streams increases the ecosystem service provision’ is based on LoE3. In addition they valued the ecosystem services, which is a subquestion of the study (‘What is the value of ecosystem services provided by streams?’). It can also be assessed for their evidence, which is especially important to guarantee multiple lines of evidence.

A second example is the governance-related question by Entenmann and Schmitt (2013): ‘Do stakeholders relate REDD+ to biodiversity conservation?’ They found that synergies between REDD+ and biodiversity conservation were assumed by stakeholders. It is an observational design (LoE_4_), receiving only 10 of 20 quality points and therefore downgraded to LoE_5_.

The third example was a systematic review of Bowler *et al*. (2010), conducted according to the guidelines of the Collaboration for Environmental Evidence (2013). They investigated the effect of greening urban areas on the air temperature to mitigate heat exposure, a management-related question. They found that green space in an urban area is on average 1°C cooler, than a built-up site. According to the quality assessment the study achieved 24 out of 26 points (92%) and it therefore remained on the originally assigned highest LoE1a.

## Common criticisms

Evidence-based practice (EBP) has faced criticism that we do not want to ignore. In the following, we discuss the most common arguments raised in evidence-based medicine and conservation (Straus and McAlister, 2000; Mullen and Streiner, 2004; Adams and Sandbrook, 2013).

### 1. Cookbook problem

*EBP is a cookbook approach denigrating professional expertise and replacing it with manualized procedures.* Best practice guidelines can not replace expertise of practitioners and best practice recommendations will highly profit of *additional* expertise, determining whether the evidence is applicable to a particular problem, bearing in mind unique circumstances (Mullen and Streiner, 2004).

### 2. EBP ignores individual variability

EBP oversimplifies complex relations and denigrates individual variability (Sackett *et al*., 1996; Feinstein and Horwitz, 1997; Straus and McAlister, 2000; Gabbay and May, 2004; Mullen and Streiner, 2004). Individual variability may overhelm general patterns, making predictions useless. However, decision-making requires the identification of general patterns to predict an outcome. Predictions based on highest available evidence provide a higher probability to reach the desired outcome and are therefore better than any unproven alternative (Mullen and Streiner, 2004).

### 3. EPB ignores qualitative data

*EBP was accused to neglect qualitative data, such as local and indigenous knowledge (Adams and Sandbrook, 2013)*. Quantitative data allow for more sensitive statistical testing and provide more information than categorical knowledge. However, qualitative data are much better than none at all and can add valuable information (Sale *et al*., 2002). As Haddaway and Pullin (2013) point out: all evidence counts. All information contribute to systematic reviews to ascertain completeness.

### 4. No evidence that EBP works

*There is insufficient evidence that EBP works better than conventional approaches (Mullen and Streiner, 2004).* EBP emerged from conventional practice over many years. Hence, there is no easy distinction between ‘the conventional approach’ and the evidence-based concept. Studies based on controlled or descriptive designs are sound scientific practice for centuries, and evidence-based research only emphasises to identify them as such. Still, we agree that the same rigour of reasoning should be applied, at a meta-level, to the concept of evidence, too. To date, too few data seem to exist to compare evidence-based decision-making with its more conventional cousin.

### 5. Environmental science is too complex for EBP

EBP works in medicine, but can not work in environmental science, because the socio-ecological system is more complex than a human body (Adams and Sandbrook, 2013). Complexity is not, in itself, a reason to abandon evidence. While certainly the medical research field is different from environmental studies, few physicians would agree that it is less complex. More importantly, however, the medical professional has typically hundreds to thousands of cases to learn from over a lifetime, while conservation ecologists work on only a very few cases. Thus, the setting for learning from experience is very different and would actually demand a more evidence-based approach to the more complex system (Gilovich *et al*., 2002).

### 6. Time and resources demanding

*EBP requires a long time to conduct a systematic review.* While in general true, this argument is misleading (Straus and McAlister, 2000). As soon as a database with systematic reviews and best-practice guidelines exists (see e.g. the Cochrane Collaboration and the Collaboration for Environmental Evidence), practitioners take less time to find an answer to their question than before.

There is further criticisms specifically addressing meta-analyses and its methodological implementation (Thompson and Pocock, 1991; Bateman and Jones, 2003). We will not elaborate on methodological details, but understand that it is crucial to properly conduct and interpret meta-analysis results and refer to (Borenstein *et al*., 2009, ch.43) for a detailed discussion of these aspects.

## Relevance for different user groups

In the previous section we have elaborated *how* to employ the evidence-based concept. Now we want to provide a few notes on *who* should use it:

1. **Scientists conducting their own studies** have to be aware how to achieve the highest possible evidence, particularly during the planning phase. Choosing a study design that provides a good evidence and respects quality aspects will substantially increase the potential contribution to our knowledge.
2. **Scientists advising decision-makers** should be aware of the evidence of information they include in their recommendations. Weighting all scientific information equally, or subjectively, runs the risk of overconfidence and bias.
3. **Decision-makers** receiving information from scientists should demand a level-of-evidence statement for the information provided, or should judge themselves the reliability having in mind the evidence-based concept.
4. We further would like to encourage **consortia, international panels and learned societies, such as the Intergovernmental Platform on Biodiversity & Ecosystem Services (IPBES), EU projects or Ecological Societies (BES, ESA, INTECOL)** to develop guidelines with recommendations on methods to best quantify, value, manage or govern a desired ecosystem service or bundle of services. This would give decision-makers a toolbox, making the common procedure (‘decision-makers seeking advice from individual scientists’) superfluous. These ‘best practice guides’ ideally exist for every single and for the sum of ecosystem services in every facet and in every ecosystem. For example we may want to ask what is the best way to quantify recreation, to value recreation, to manage recreation and to use governance strategies that fosters sustainable recreation in a temperate forest. Each best practice guide would clearly state its level of evidence. At a higher level, where the sum of all ecosystem services in one ecosystem need to be evaluated, it would make sense to have a best practice guide on how to measure, say, the total (economic) value (e.g. summing individual values up with a strategy to avoid double-counting (Boyd and Banzhaf, 2007; DEFRA, 2007)). All this may sound unrealistic, given the huge number of methods, ecosystem services, management and governance options and so forth. However, in medicine, national and international learned societies set up assessment and guideline boards for exactly this purpose (often with governmental support, e.g. the UK’s National Institute for Health and Care Excellence (NICE) www.nice.org.uk or Germany’s IQWiG www.Iqwig.de). There are currently 261 recognised diseases with over 12000 sub-categories (ICD-10). This is certainly at the same scale as the challenges faced by ecosystem service science.

## Conclusion

We introduced the evidence-based concept in ecosystem service science, encompassing a scale to judge the available evidence and a quality checklist to facilitate critical appraisal. We further showed in detail and illustrated with examples how to use the concept. Additional support and guidance can be obtained by the Collaboration of Environmental Evidence (www.environmentalevidence.org).

The evidence-based ecosystem service science does not suggest a specific management strategy. It is by no mean a contradiction or replacement to adaptive management or other management concepts. Rather, it complements these approaches, emphasising that whatever is used should be used with the awareness of how approved our knowledge is.

Wrong decisions can have strong negative consequences. This is particulary painful, if studies providing high evidence were available, but instead decisions were based on myth or low evidence studies. Taking again an example from medicine, child mortality from sudden child death was unnecessary high for decades due to wrong recommendations based on low evidence, ignoring the higher evidence available (Gilbert *et al*., 2005). Especially on topics whith various and contradicting opinions, it is important to continuously summarise and update the available evidence. If farmers have no reliable information on the management of natural pest control versus pesticides (Wright *et al*., 2013), their actions may result in huge and avoidable economic loss or even directly affect human health.

It should have become clear that evidence-based ecosystem service science concerns scientists as well as decision-makers and the general public. In the interest of a responsible use of environmental resources and processes, we strongly encourage embracing evidence-based practice as paradigm for all research contributing to ecosystem service.

## Acknowledgements

This work was supported by the 7th framework programme of the European Commission in the project ‘Operational Potential of Ecosystem Research Applications’ (OPERAs, grant number 308393).

