## Supplementary Material for "A plea for evidence in ecosystem service science: a framework and its application"

Anne-Christine Mupepele & Carsten F. Dormann
University of Freiburg

We applied the evidence-based concept to 12 case studies, 3 of them already mentioned in the main document. The details about the context, facet, the design as well as the critical appraisal (the quality assessment) are given in Table S1. We further present the quality checklist that was used to determine the study quality in all 12 case studies (Table S2). We covered a broad range of case studies, providing an example for all facets (quantification, valuation, management and governance) and all different study designs.

Table S1: Studies ranked according to the evidence-based approach. The key columns (grey section) are the question, the outcome and the final level of evidence (LoE). They are supported by context, facet and the evidence assessment with study design and quality assessment (critical appraisal).

| | 1. Question, outcome and the context | | | | 2. Evidence Assessment | | | | |
|---|---|---|---|---|---|---|---|---|---|
| Reference | Context: Ecosystem services; Ecosystem(s); Location | Facet | Question investigated | Outcome | Final level of evidence | 2a. Study design -> | Level of evidence | 2b.Quality points (see checklist S2) -> | Quality score |
| Bowler et al. 2010 | Air conditioning in urban space; cities; global | Management | Can human exposure to heat be mitigated by greening urban spaces | Green space in an urban area is on average 1°C cooler, than a built-up site. | LoE1 | Systematic review | LoE1a | 24/26 | 0.92 |
| Lindhjem 2007 | Non-timber forest ecosystem services, mainly recreation; forests; Norway, Sweden, Finland | Valuation | Review of people's willingness to pay for non-timber forest ecosystem services and explain systematic variation | Willingness to pay (WTP) is insensitive to forest size; WTP is higher if people are asked as individuals than on behalf of their households | LoE2 | Conventional review | LoE1b | 20/27 | 0.74 |
| Ayanu et al 2012 | Crops and biomass production, above ground carbon storage, amount of pollutants removed from the air, soil retained, water purification, storm mitigation, pest control; all; global | Quantification | Analysing advantages and disadvantages of several remote sensing procedures | Description of various remote sensing techniques. Claim for more studies assessing validity, reliability and uncertainty of remote sensing procedures in quanitiying and mapping ecosystem services | LoE2 | Conventional review | LoE1b | 10/18 | 0.56 |
| Liu et al. 2008 | Timber, soil erosion, carbon sequestration, recreation through wildlife observing; forests; China | Governance | Effects of Payment for Ecosystem Services (PES) - What is the socioeconomical and ecological impact of two in China implemented PES programs? | Socialeconomical impact: positive&negative, e.g. total income increased, decline in local revenues for local governments. Ecological benefits: Timber harvest decreased but timber import increased. Carbon sequestration increased, area suffering from soil erosion declined. | LoE4 | Conventional review | LoE1b | 5/20 | 0.25 |
| Millar et al. 2010 | Soil erosion protection; grassland; USA | Quantification | Investigating the effect of sod farming on soil loss. | Net loss of productive soil under sod farming is larger than the tolerable soil loss. | LoE3 | Case-control | LoE2 | 14/25 | 0.56 |
| Acuna et al. 2013 | Food (fish), retention of organic and inorganic matter; river,forests; Iberian Peninsula | Management | How does adding dead wood to stream channels affect the provision of selected ecosystem services? | Restoration of natural wood loading in streams greatly increases the ecosystem services they provide. | LoE3 | Before-after control-impact | LoE2 | 14/24 | 0.58 |
| Lara et al. 2009 | Food (fish); marine; mediterranean | Quantification | Developping an index that estimates fish density, biomass and production in dependance of environmental variables | Index etimating fish density, biomass and production in dependance of surface area/volume ratio, water volume with sufficient oxygen, conductivity, cholorphyll a concentration perimeter | LoE4 | Correlative | LoE3 | 10/19 | 0.53 |
| Barkmann et al. 2008 | Fibre, water, recreation/biodiversity, cacao; agroforestry; Indonesia | Valuation | What is the value of the central ecosystem services the hydrological ecosystem provides? | Willingness to pay for improving water availability equals about 1% of mean cash income of households. | LoE5 | Regression model | LoE3 | 10/20 | 0.50 |
| Xie et al. 2011 | Improved air quality; city; China | Quantification | Quantification of $CO_2$ sequestration, $O_2$ production, dust removal of different plant species | Absolute numbers for $CO_2$ sequestration, $O_2$ production and dust removal and the differences between plant species | LoE5 | Descriptive | LoE4 | 8/18 | 0.44 |
| Karimzadegan et al. 2007 | gas regulation, pollination, pest control and others; forests; Iran | Valuation | What is the value of Irans forests and rangeland ecosystem services | A value [$] for Irans forests and rangeland | LoE5 | Descriptive | LoE4 | 8/21 | 0.38 |
| Entenmann and Schmitt 2013 | Biodiversity; forests; Peru | Governance | Do stakeholders relate REDD+ to biodiversity conservation? | Yes, synergies between REDD+ and biodiversity conservation were assumed. | LoE5 | Descriptive | LoE4 | 10/20 | 0.50 |
| Desanker 2005 | climate stabilisation; all; Africa | Governance | How can the Clean Development Mechanism better engaged in Africa? | Projects should be initiated by locals rather than esternal groups, additional CDM fund - not only carbon money, values from forest products and services should be factores into the economic Analysis | LoE5 | Expert opinion | LoE5 | not required, it is already on the lowest level of evidence | |

### Table S2: **Quality checklist** used to obtain the quality points (2b in Table S1)

| | | Reference: | Bowler et al. 2010 | Lindhjem 2007 | Ayanu et al 2012 | Liu et al. 2008 | Millar et al. 2010 | Acuna et al. 2013 | Lara et al. 2009 | Barkmann et al. 2008 | Xie et al. 2011 | Karimzadegan et al. 2007 | Entenmann and Schmitt 2013 | Desanker 2005 |
|---|---|---|---|---|---|---|---|---|---|---|---|---|---|---|
| | 1 | **Correspondence (Does the question match the answer?)** | **yes** | **yes** | **yes** | **yes** | **yes** | **yes** | **yes** | **yes** | **yes** | **yes** | **yes** | |
| | 2 | Are the assumption used in the study reasonable? | yes | yes | yes | / | yes | yes | yes | yes | yes | yes | no | |
| | 3 | Internal validity: Do design and implementation avoid a high risk of bias? | yes | yes | no | no | yes | yes | no | yes | no | no | no | |
| | 4 | External validity/relevance: Is the result transferable to other scenarios with the same context? | yes | no | yes | yes | yes | yes | yes | no | yes | yes | yes | |
| | 5 | Are multiple lines of evidence considered? | no | no | no | no | no | no | no | / | no | no | no | |
| Data collection | 6 | Was the target population/area defined in space, time and size? | yes | yes | no | yes | yes | no | yes | no | no | yes | yes | |
| Data collection | 7 | Was a sampling population/area defined? (Which population/area was sampled?) | yes | yes | no | no | yes | yes | yes | yes | yes | yes | yes | |
| Data collection | 8 | Were potential differences between the target population and the sampling population considered? | yes | no | / | / | no | / | no | / | / | no | no | |
| Data collection | 9 | Were the methods described in sufficient detail to permit replication? | yes | yes | / | no | yes | yes | yes | no | yes | yes | yes | |
| Data collection | 10 | Was the sample size appropriate? | yes | yes | yes | no | yes | yes | no | yes | no | no | yes | |
| Data collection | 11 | **Was probability sampling used for constructing the sample?** | **/** | **/** | **/** | **/** | **no** | **no** | **no** | **no** | **no** | **no** | **no** | |
| Data collection | 12 | If secondary data are used, did an evaluation of the original data take place? | yes | yes | yes | no | / | / | / | / | / | no | / | |
| Analysis | 13 | Is the choice of statistical/analytical method justified and comprehensively explained? | yes | yes | / | / | yes | yes | yes | yes | no | / | yes | |
| Analysis | 14 | Are variables and statistical measures given? | yes | yes | / | / | yes | no | yes | yes | yes | no | yes | |
| Analysis | 15 | **Was accuracy/uncertainty assessed and reported?** | **yes** | **yes** | **/** | **no** | **no** | **yes** | **no** | **no** | **no** | **no** | **no** | |
| Results | 16 | Are results consistent and homogeneous? | yes | yes | / | yes | yes | no | yes | yes | yes | yes | yes | |
| Results | 17 | Magnitude of effect: Is the effect large (and without large uncertainty) | no | no | / | | yes | no | no | / | no | no | no | |
| Results | 18 | Attrition bias: Are non-response/drop-outs given and is their impact discussed | yes | / | no | no | / | / | / | no | / | / | no | |
| | | Design-dependent aspects: | | | | | | | | | | | | |
| Review | 19 | Low probability of publication bias? E.g. results reporting a negative relationship were probably not included | yes | yes | no | no | / | / | / | / | / | | / | |
| Review | 20 | **Is the review based on high evidence individual studies (several level 2 or level 3 studies)?** | **yes** | **no** | **yes** | **no** | **/** | **/** | **/** | **/** | **/** | | **/** | |
| Review | 21 | Validity – Do the studies included respond to the same question? | yes | yes | yes | / | / | / | / | / | / | | / | |
| Review | 22 | Was the literature searched in a systematic way? | yes | no | yes | no | / | | / | | | | / | |
| Review | 24 | Was a meta-analysis (in the strict sense: see Borenstein et al. 2009) indcluded? | yes | yes | no | no | | | | | | | | |
| Review | 24 | Were any other quantitative summary statistics provided? | yes | yes | no | no | | | | | | | | |
| Studies with a reference | 25 | **Selection bias: Was the assignment of case-control groups randomized?** | **/** | **/** | **/** | **/** | **no** | **no** | **/** | | **/** | | **/** | |
| Studies with a reference | 26 | **Were groups designed equally, aside from the investigated point of interest?** | **/** | **/** | **/** | **/** | **no** | **yes** | **/** | **/** | **/** | | **/** | |
| Studies with a reference | 27 | Performance bias: Was the sampling blinded, e.g. researchers taking samples of a specific area wouldn't know which differences are between these areas? | / | / | / | / | no | no | / | / | / | | / | |
| Studies with a reference | 28 | Were there sufficient replicates of treatment and reference groups? | / | / | / | / | yes | yes | / | / | / | | / | |
| | | Category-dependent aspects: | | | | | | | | | | | | |
| Valuation | 29 | Were future values of ecosystem services considered? | / | yes | / | / | / | / | / | no | / | no | / | |
| Valuation | 30 | If future values were considered, were they discounted with a well-motivated discount rate? | / | yes | / | / | / | / | / | / | / | / | / | |
| Valuation | 31 | If aggregate economic values for a population were estimated, was this estimation consistent with the sampling procedure and the definition of the population? | / | / | / | / | / | / | / | no | / | no | / | |
| Valuation | 32 | If valuation took place in form of a questionnaire, was the study pre-tested and piloted? | / | / | / | / | / | / | / | yes | / | / | / | |
| | | Quality points | 24/26 | 20/27 | 10/18 | 5/20 | 14/25 | 14/24 | 10/19 | 10/20 | 8/18 | 8/21 | 10/20 | |
